# A method for separating bacterial cultures into single cells and defined size fractions of biofilm aggregates

**DOI:** 10.1101/2024.01.25.577188

**Authors:** Regitze Lund Nielsen, Thomas Bjarnsholt, Tim Holm Jakobsen, Mads Lichtenberg

**Affiliations:** Costerton Biofilm Center, Department of Immunology and Microbiology, University of Copenhagen, Denmark; Department of Clinical Microbiology, Copenhagen University Hospital, Denmark

**Author notes:** Corresponding author, Blegdamsvej 3B, 2200 Copenhagen, Denmark.

## Abstract

*In vitro* microbiological experiments that aim to describe differences between planktonic and biofilm aggregate populations are challenging since liquid batch cultures contain a mix of both. Here, we present a simple method for fractioning a bacterial liquid batch culture into aggregates and single cells.

Stackable cell strainers with mesh sizes of 30 μm and 10 μm were used to filtrate 6 day old batch cultures of *Pseudomonas aeruginosa* to produce size fractions of 0-10μm and >30μm. By confocal laser scanning microscopy measurements, we show that 95.5% of the total biomass was <10 μm in the “0-10μm size fraction” and that 92.5% of the total biomass was >30μm in the “>30μm size fraction”.

Furthermore, the adjustment of bacterial concentration using CFU/ml was validated by quantifying the total DNA of viable bacteria in the two size fractions after DNase treatment to deplete eDNA and DNA from dead bacteria. Surprisingly, this showed that adjusting the bacterial concentration using CFU/ml was a valid method with no significant differences in total DNA from viable bacteria.

## Introduction

Decades of research have shown that bacteria in biofilms behave differently than they do when living as planktonic or freely suspended cells. Especially, surface attached biofilms have received a lot of attention due to their conspicuous nature and ease of handling (1). However, non-attached aggregates are receiving increasing attention and have been shown to possess many pheno- and genotypic similarities to surface biofilm although important differences exist (2).

Studies have shown that liquid batch cultures contain a mix of single cells and aggregates (3, 4) and the presence of aggregates can influence the outcome of experiments (5, 6). Thus, using liquid batch cultures to start experiments that aim at investigating differences between planktonic bacteria and aggregates are not ideal as the culture will contain a mixture of both. Alternative approaches could entail harvesting aggregates from a biofilm and comparing them to a young exponential phase culture that mostly consist of single cells, but using this approach leaves little control over the size of aggregates. Additionally, one would compare two populations displaying very different metabolic states which can also have profound impact on the outcome of in vitro experiments (7, 8).

Here, we aim at developing a simple method for isolating aggregates and single cells from the same culture, allowing precise control of the size of aggregates as well as having bacteria isolated from the same growth conditions.

## Materials and Methods

### Bacterial strains and growth conditions

*P. aeruginosa* isolates PAO1 (PAO0001) obtained from the Pseudomonas Genetic Stock Center and PAO1 tagged with a green fluorescent protein (GFP) expressed on plasmid pMRP9(9) were used in this study. To select bacteria expressing the GFP-encoded plasmid, a fluorescent microscope was used to collect colonies containing fluorescent bacteria of PAO1. Freezer stock (-80°C) of PAO1 and GFP-tagged PAO1 was streaked on a Lysogeny-Broth (LB) agar plates (Panum Substrate Department, Copenhagen, Denmark) and incubated overnight at 37°C. The following day colonies were collected from the plates and inoculated in 100 ml LB media (Panum Substrate Department, Copenhagen, Denmark) supplemented with 0.3% glucose (Panum Substrate Department, Copenhagen, Denmark) in 250 ml Erlenmeyer flasks. The cultures were incubated at 37°C on a shaker set to 180 revolutions per minute (RPM) for six days.

### Fractionation of bacterial culture into planktonic cells and aggregates

After six days of incubation of the bacterial culture, 45ml was poured over a stack of cell strainers (PluriSelect, Leipzig, Germany) with mesh sizes of 30 μm and 10 μm (Fig. 1). Aggregates collected in the 30 μm strainer were isolated and collected by carefully scraping and backwashing the filter, and the planktonic cells were collected in the filtrated liquid. The fraction containing aggregates was washed twice with saline to reduce the number of planktonic cells. The wash was done by pipetting off the upper part of the fraction after the aggregates were allowed to settle, as done previously (6). The volume removed during the wash was replenished with saline. The size fractions were kept at 4°C to prevent further growth until CFU/ml was determined.

**Figure 1:**
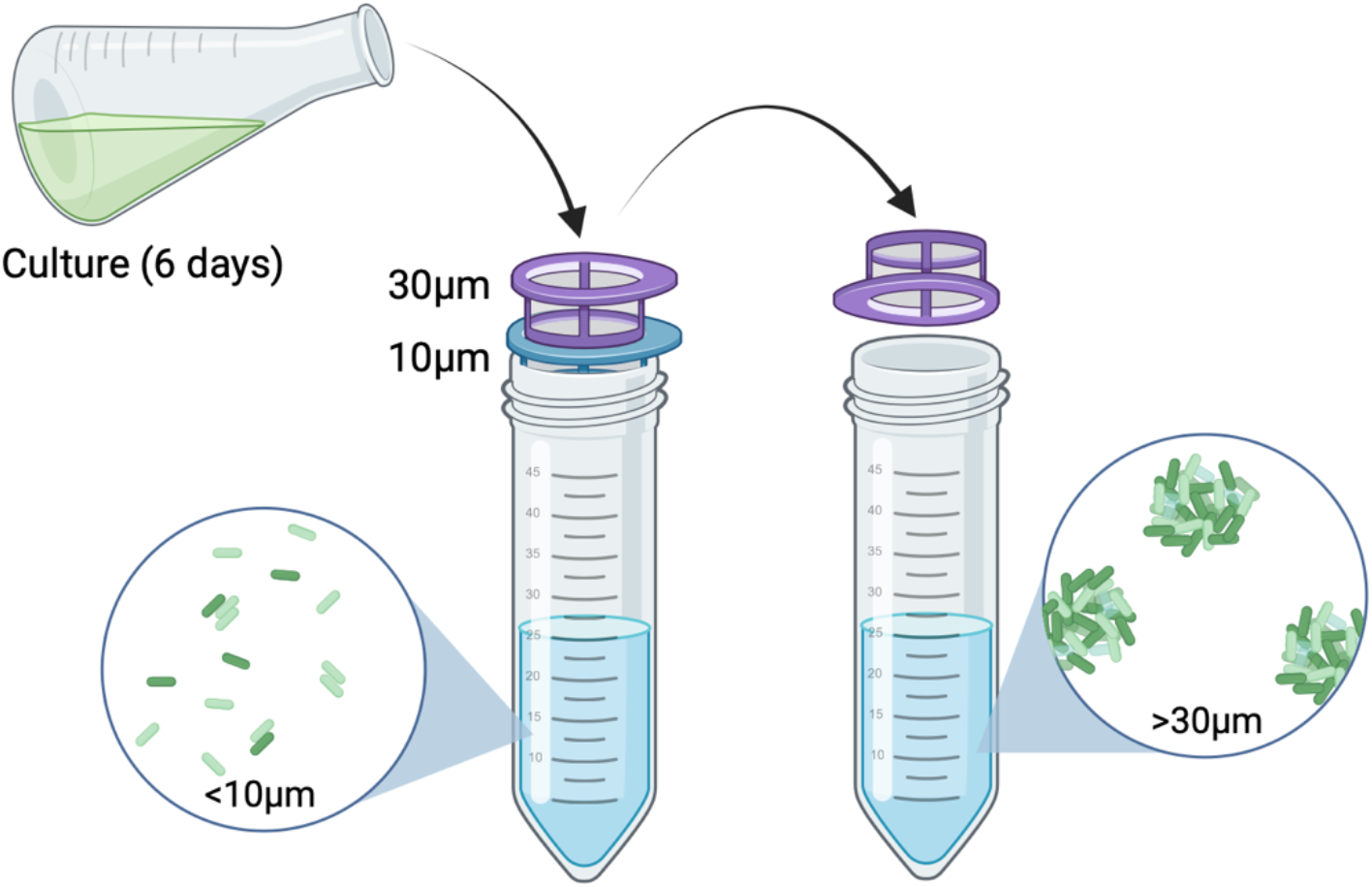
Graphical representation of the filtration method used to separate single cells (0-10 μm) from aggregates >30μm.

### Enumeration of bacteria

Colony Forming Units (CFU) were used to determine the number of viable cells in the different size fractions. 10-fold dilution series were made in an appropriate number of dilutions by transferring 100 μl of the bacteria-containing solution to a Widal tube containing 900 μl saline (0.9%) accompanied by thorough vortexing before continuing the dilution row. In three technical replicates, 10 μl of the different dilutions were spotted on LB agar plates (Panum substrate department, Denmark, Copenhagen). The LB plates were incubated at 37°C until the following day, when colonies were counted at a dilution with an appropriate number of colonies per spot (4-40 colonies). The number of colonies, the dilution, and the volume of the sample spotted, were used to determine CFU/ml.

### DNA extraction of planktonic cells and aggregates

To verify whether CFU/ml was an appropriate measure to compare the number of viable bacteria in aggregates and planktonic cells, DNA extraction of the fractions adjusted to concentrations of 10^6^, 10^7^, and 10^8^ CFU/ml was carried out. The cells were first treated with DNase I (ZYMO Research, Irvine, California, USA) to remove extracellular DNA and DNA from dead cells. Then the DNA from viable bacteria was extracted using a blood and tissue extraction kit (Qiagen, Hilden, Germany).

1 ml of cells with different CFU/ml were centrifuged at 10,000 x G for 10 minutes, and the supernatant was discarded. The pellet was resuspended in 200 μl saline with 10 μl of DNase I with a concentration of 1U/ng and 10 μl DNase buffer, which then was allowed to incubate for 10 minutes at 37°C on a shaker set to 100 rpm. The enzyme was inactivated by transferring the samples to a heating block set to 65°C for 10 minutes. The suspension was centrifuged for 10 minutes at 10,000 x G, the pellet was resuspended in 200 μl saline, and the supernatant was discarded. The cells were washed twice by repeating the following steps two times: After centrifugation, the pellet was resuspended in 200 μl saline, and the supernatant was discarded. The suspension was centrifuged for 10 minutes at 10,000 x G.

The rest of the DNA extraction was carried out according to the manufacturer’s instructions for the blood and tissue extraction kit (Qiagen, Hilden, Germany). Finally, the DNA concentration was determined with a Qbit 4 fluorometer (Thermo Fisher Scientific Inc., Waltham, Massachusetts, USA), by incubating 20 μl of the sample in 180 μl dsDNA 1x solution (Thermo Fisher Scientific Inc., Waltham, Massachusetts, USA).

### Confocal laser scanning microscopy

A fractionated PAO1 culture was diluted to 10^6^, 10^7^, and 10^8^ CFU/ml for aggregates and planktonic cells, respectively. The fractions were stained with Syto9 (Merck, Darmstadt, Germany) with a final concentration of 2.5μM and added to channel slides (uncoated μ-slide VI-flat, Ibidi, Gräfelfing, Germany). The samples were analysed with a confocal laser scanning microscope, CLSM 880 (Zeiss, Oberkochen, Germany), using a 20x/0.8 M27 objective and a frame size of the images at 425x425 μm. An Argon laser with an excitation wavelength of 488 nm was used to excite the fluorophore. The microscope was set to detect emitted light for wavelengths ranging from 494-561 nm. Surface areas of the bacteria in the images were quantified using Imaris (version x64 9.7.2), and areas were grouped into intervals of diameters.

## Results and discussion

A method was developed to separate planktonic and aggregated bacteria for use in later experiments. Confocal microscopy images indicated that the planktonic fraction primarily consisted of planktonic cells, while the fraction containing aggregates mainly consisted of aggregates (Fig. 2A).

**Figure 2:**
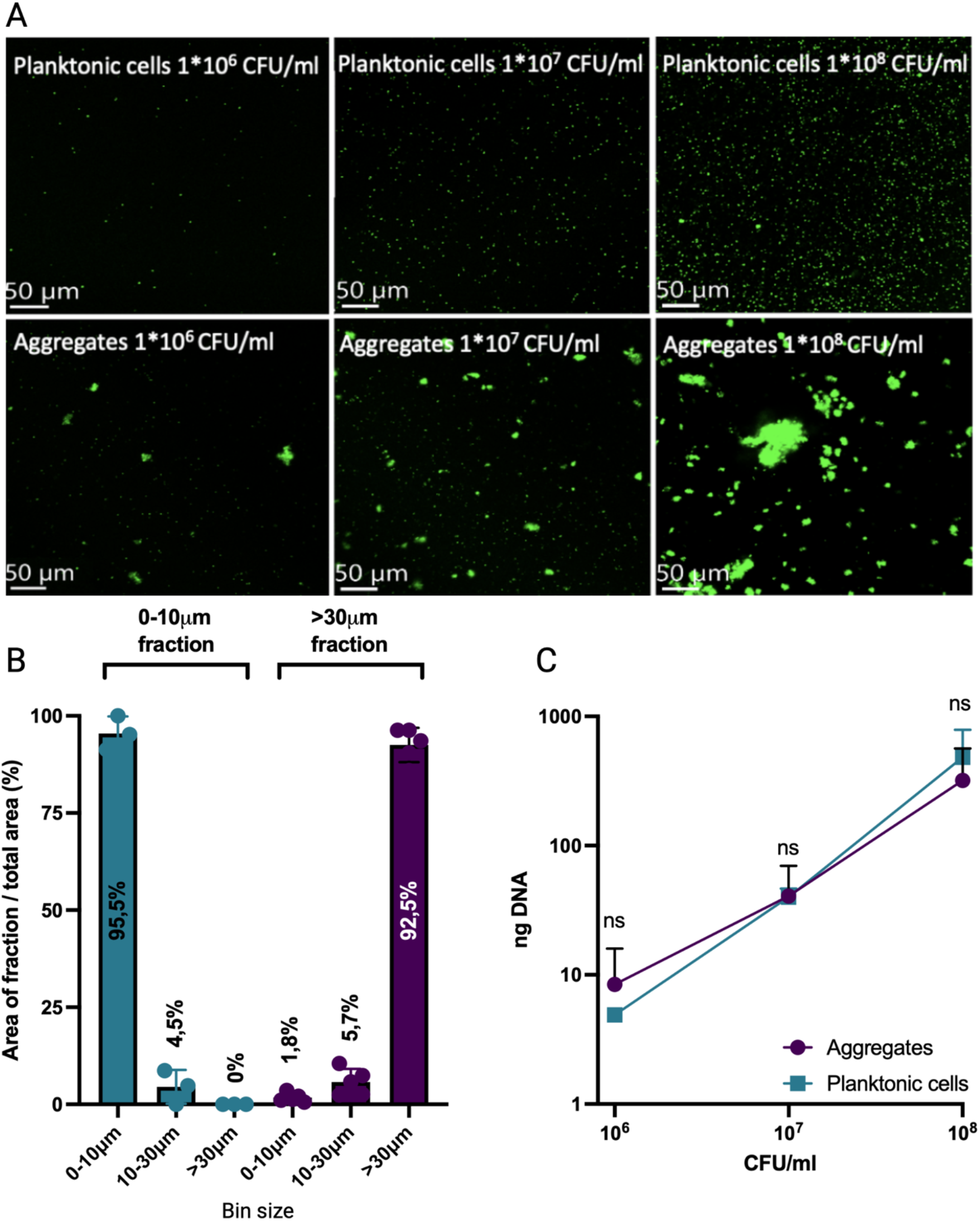
Fractionated bacterial cultures into planktonic cells and aggregates. **A)** Visualization of biomass of the planktonic and aggregate fractions diluted to different CFU/ml (representative images of three individual experiments). **B)** Biomass distribution within non-diluted fractions (Mean±SD of 5 technical replicates, from each 3 biological replicates of the planktonic fraction and 5 of the aggregated fraction). **C)** DNA (in ng) extracted from planktonic cells (blue) and aggregates (purple) adjusted to different CFU/ml. Data was analyzed by 2way ANOVA. Datapoints represent means±SD; n=3.

Quantification of the biomass (by estimating surface areas) of the planktonic (0-10μm) and aggregate (>30μm) fractions confirmed that 95.5% of the total biomass of the planktonic fraction had a diameter between 0 and 10μm while the remaining 4.5% was bound in aggregates of 10-30μm (Fig. 2B). Thus, no aggregates >30 μm was observed in this fraction. The aggregate fraction had 92.5% of the total biomass with a diameter >30μm while 5.7% was bound in aggregates of 10-30μm and only 1.8% was found as aggregates <10μm (Fig. 2B). Thus, the 0-10 μm fraction was completely free of large aggregates while the aggregate fraction >30μm was highly enriched for large aggregates while only having a minor proportion of smaller aggregates and/or single cells. It was expected that the >30 μm fraction would contain a small proportion of aggregates <30μm since some will adhere to the filter and be transferred during washing. Similarly, breaking up of larger aggregates during washing and pouring will result in smaller aggregates than intended. However, we show that the majority of the biomass was bound in the intended size fractions.

Enumeration of bacteria in non-attached aggregates and biofilms have traditionally been considered unreliable using plate count method since an aggregate of thousands of bacteria will still resemble one colony forming unit and therefore the true number of bacteria can be grossly underestimated. Therefore, we attempted to validate if the CFU adjusted fractions contained the expected number of bacteria. We did this by quantifying the DNA content of viable bacteria by first removing extracellular DNA and DNA from dead bacteria by treating with DNase I and washing several times. DNA content in the viable planktonic and aggregate fractions was then extracted to verify the quantification of bacteria by CFU counting. We found that the amount of DNA between the planktonic and aggregate fractions diluted to 10^6^, 10^7^, and 10^8^ CFU/ml, respectively, were not statistically different (Fig. 2C). Unexpectedly, this suggests that, despite small differences, comparable numbers of viable bacteria were present in the two fractions adjusted to the same CFU/ml. We speculate that aggregates could contain a large proportion of dead bacteria and therefore the aforementioned limitation in the plate count method is counterbalanced by a lower number of viable bacteria than expected from the volume of aggregates. The presence of dead bacteria was also confirmed by CLSM observations using live/dead stain but was not further quantified here.

*In vitro* bacteriological experiments are highly sensitive to variations in the inoculum and we have previously shown that it can influence the interpretation and outcomes of experiments (5, 6, 8, 10). Therefore, the finding that the plate count method can actually be used to adjust cultures of planktonic and aggregated bacteria to the same concentration of viable bacteria warrants further validation. But it seems at least for the conditions used in this study, the method is applicable.

In summary, we present a simple method for separating a bacterial culture into aggregates and single cells using accessible and disposable equipment that do not require specialized setups and thus is able to be used widely.

